# Rapid genotyping of porcine reproductive and respiratory syndrome virus (PRRSV) using MinION nanopore sequencing

**DOI:** 10.1101/2023.02.23.529665

**Authors:** Leonardo Cardia Caserta, Jianqiang Zhang, Pablo Piñeyro, Diego G. Diel

## Abstract

The global distribution and constant evolution are challenges for the control of porcine reproductive and respiratory syndrome virus (PRRSV), one of the most important viruses affecting swine worldwide. Effective control of PRRSV benefits from genotyping, which currently relies on Sanger sequencing. Here we developed and optimized procedures for real-time genotyping and whole genome sequencing of PRRSV directly from clinical samples based on targeted amplicon- and long amplicon tiling sequencing using the MinION Oxford Nanopore platform. Procedures were developed and tested on 154 clinical samples (including lung, serum, oral fluid and processing fluid) with RT-PCR Ct values ranging from 15 to 35. The targeted amplicon sequencing (TAS) approach was developed to obtain sequences of the complete ORF5 (main target gene for PRRSV genotyping) and partial ORF4 and ORF6 sequences of both PRRSV-1 and PRRSV-2 species. After only 5 min of sequencing, PRRSV consensus sequences with identities to reference sequences above 99% were obtained, enabling rapid identification and genotyping of clinical PRRSV samples into lineages 1, 5 and 8. The long amplicon tiling sequencing (LATS) approach targets type 2 PRRSV, the most prevalent viral species in the U.S. and China. Complete PRRSV genomes were obtained within the first hour of sequencing for samples with Ct values below 24.9. Ninety-two whole genome sequences were obtained using the LATS procedure. Fifty out of 60 sera (83.3%) and 18 out of 20 lung samples (90%) had at least 80% of genome covered at a minimum of 20X sequence depth per position. The procedures developed and optimized in this study here are valuable tools with potential for field application during PRRSV elimination programs.

## Introduction

Porcine reproductive and respiratory syndrome virus (PRRSV) is a single-stranded positive sense RNA virus with a genome of approximately 15kb in length classified within the family *Arteriviridae*. Currently, PRRSV is classified into two species: *Betaarterivirus suid 1* (PRRSV-1, also referred to as European PRRSV) and *Betaarterivirus suid 2* (PRRSV-2, North American PRRSV) (1). PRRSV is one of the most economically significant swine pathogens worldwide. The infection can lead to reproductive failure, clinical disease in sows and boars, neonatal losses, and respiratory disease in piglets. The estimated annual cost of PRRSV for the U.S. pork industry is around 664 million dollars (2).

Traditional PRRSV diagnostic relies on real-time reverse transcriptase PCR (RT-PCR) detection of the virus in serum, oral fluids, semen and/or tissue samples (3). Given the high genetic diversity of PRRSV and the broad use of modified live virus (MLV) vaccines, PRRSV detection by RT-PCR is usually complemented by sequencing ORF5 gene encoding the glycoprotein 5 (GP5) followed by restriction fragment length polymorphism (RFLP) analysis. The predicted ORF5 cut pattern of three specific restrictions enzymes (MluI, HincII, and SacII) is used to classify PRRSV into specific RFLP profiles (i.e. 1-8-4, 1-7-4, etc.) (4). While the method enabled important discoveries and advances related to PRRSV for more than a decade, this approach only provides limited information on the genetic makeup of the virus.

Sequencing of ORF5 allowed the classification of PRRSV-2 viruses into nine lineages: 1 to 9 (5). Lineage 9 was the most prevalent from 2009 to 2010, but its frequency decreased coinciding with the emergence or re-emergence of lineage 1 as the dominant lineage, which reached a prevalence of ~90% by 2015. Viruses within lineages 1, 5, 7, and 8 are used in commercial vaccines in the United States with the vaccine belonging to lineage 5 being the most widely used historically (6).

Rapid evolution and the genetic diversity of the virus are major issues affecting disease control and management. The emergence of highly pathogenic PRRSV strains in China and the constant discovery of variant viruses underscore the need for improved diagnostics and rapid sequencing approaches that enable complete genetic characterization of the viruses circulating in the field (6–8). Up to 40% of the U.S. breeding herds experience PRRSV outbreaks every year (9), thus it is important to identify the virus that is present and define whether the outbreak(s) is/are a result of a new virus introduction(s), or of the emergence of a variant strain(s). During the fall of 2020, a PRRSV outbreak with extremely high production losses was reported in the Midwestern U.S., with the first peak occurring between October and December, and the second starting in April 2021. Results of ORF5 and whole genome sequencing suggested this outbreak was represented by the emergence of a new variant within Lineage 1C and showed low positive predictive values for either RFLP pattern 1-4-4 or Lineage 1C (10).

The use of technologies and strategies to rapidly identify PRRSV that simultaneously yield the complete genetic sequences of the virus may contribute to improved control strategies (e.g. modified live virus vaccine selection) in the field. Full-length genome sequencing and careful phylogenetic analyses are required to draw appropriate conclusions on the homology between vaccine strains and circulating field viruses (11). Whole genome sequencing is currently not routinely performed as a surveillance tool by most of the swine industry in part because of the high cost (10). The use of the so-called second-generation sequencing platforms (Roche 454, SOLiD and Illumina) enabled massive parallel sequencing generating an enormous amount of data for a reasonable price. However, some setbacks of these platforms are the requirement of expensive equipment, and challenging *de novo* assembly of the genome due to the production of short reads (highly-fragmented yield) or insufficient genome coverage (12). To overcome these hurdles, MinION sequencing has greater potential for use in the field and for simultaneous detection and real-time diagnostic identification of pathogens when compared to other third-generation sequencing technologies. The MinION platform is portable (around 100g), cheap, and can be easily connected to computers and other portable devices without the need for internet connection for data manipulation (12).

The use of the MinION platform as a genotyping tool to be applied directly in clinical samples has been reported for other RNA viruses including Ebola virus, and some arboviruses present in low concentrations such as Zika virus (13–15). The use of tiling amplicon multiplex PCR for genome enrichment has enabled the generation of hundreds of thousands of SARS-CoV-2 genomes that were made available in public databases in an unprecedented effort and speed to track the virus evolution (16).

Deep sequencing of partial genes has been described as another approach for genotyping viruses using MinION. Enterovirus, infectious bronchitis virus, Newcastle disease virus and infectious laryngotracheitis virus were successfully sequenced and identified at genotype and lineage level (21–25).

Fast genetic characterization, yielding complete or near complete PRRSV genome sequences, a highly prevalent and diverse pathogen of swine, would improve our understanding of PRRSV epidemiology and allow clinicians and field veterinarians to direct and implement prevention and control measures. In this study we present the development, optimization and use of assays for rapid genotyping of PRRSV directly from clinical samples based on partial (ORF5) and/or whole genome sequencing using the MinION platform.

## Materials and Methods

### Clinical samples and RNA extraction

Swine clinical samples originated from different herds throughout the U.S. between 2019 and 2021 were submitted to the Veterinary Diagnostic Laboratory at Iowa State University for diagnostic investigation using a PRRSV RT-PCR assay. A total of 154 PRRSV-2 PCR-positive samples were used and included: 20 lungs, 60 sera, 35 oral fluids and 39 processing fluid samples with RT-qPCR Ct values ranging from 15 to 35 (Table S1). Samples were clarified by centrifugation at 2,000 rpm for 10 minutes and subjected to nucleic acid extraction using the cador Pathogen 96 QIAcube HT Kit (Qiagen) in a QIAcube HT automated extractor (Qiagen).

### Library preparation for targeted amplicon sequencing (TAS)

For targeted amplicon sequencing (TAS), the ORF4-6 genomic region was amplified directly from the clinical samples using the SuperScript™ IV One-Step RT-PCR System (Thermo Fisher Scientific, Waltham, MA). Primers were designed using Primer3 (26) in the Geneious Prime 2019 software (https://www.geneious.com) targeting a ~1,546 bp region covering the full length ORF5 (envelope protein), and partially covering ORF4 and ORF6 (membrane protein) of PRRSV-1 and PRRSV-2 (Figure 1). Universal Oxford nanopore-compatible adapter sequences were added to the 5’ end of each primer sequence to allow PCR-based barcoding. Primer sequences are provided in Table 1. The RT-PCR reaction was carried out using 5 μl of RNA and the thermocycling conditions as follows: 50°C for 10 min; 98°C for 2 min; 40 cycles of 98°C for 10 s, 70°C for 10 s, and 72°C for 45 s, followed by one cycle at 72°C for 5 min. After purification of amplicons with AMPure XP beads (Beckman Coulter, Brea, CA) at a 1.6:1 volumetric bead-to-DNA ratio and quantification using the dsDNA High Sensitivity Assay kit on a Qubit^®^ fluorometer 3.0 (Thermo Fisher Scientific), samples were diluted to 0.5 nM in a total of 24 μl and used as input for library preparation following the PCR barcoding (96) amplicon (SQK-LSK109) protocol (Oxford Nanopore Technologies [ONT]). Final DNA libraries were loaded in a FLO-MIN106 R9.4 flow cell to start 3-hour sequencing runs. Pools of 24 or 48 barcoded samples were sequenced and the total number of reads obtained within 1 hour were analyzed.

**Figure 1.**
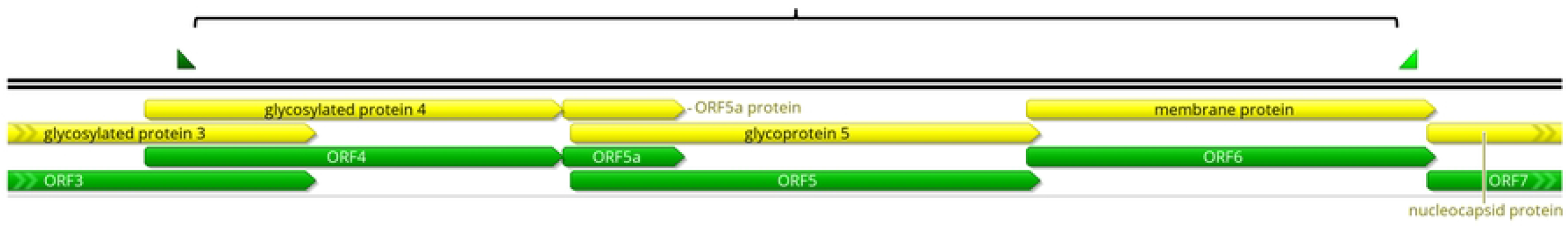
Region of PRRSV genome covered by the TAS primers.

**Table 1.**
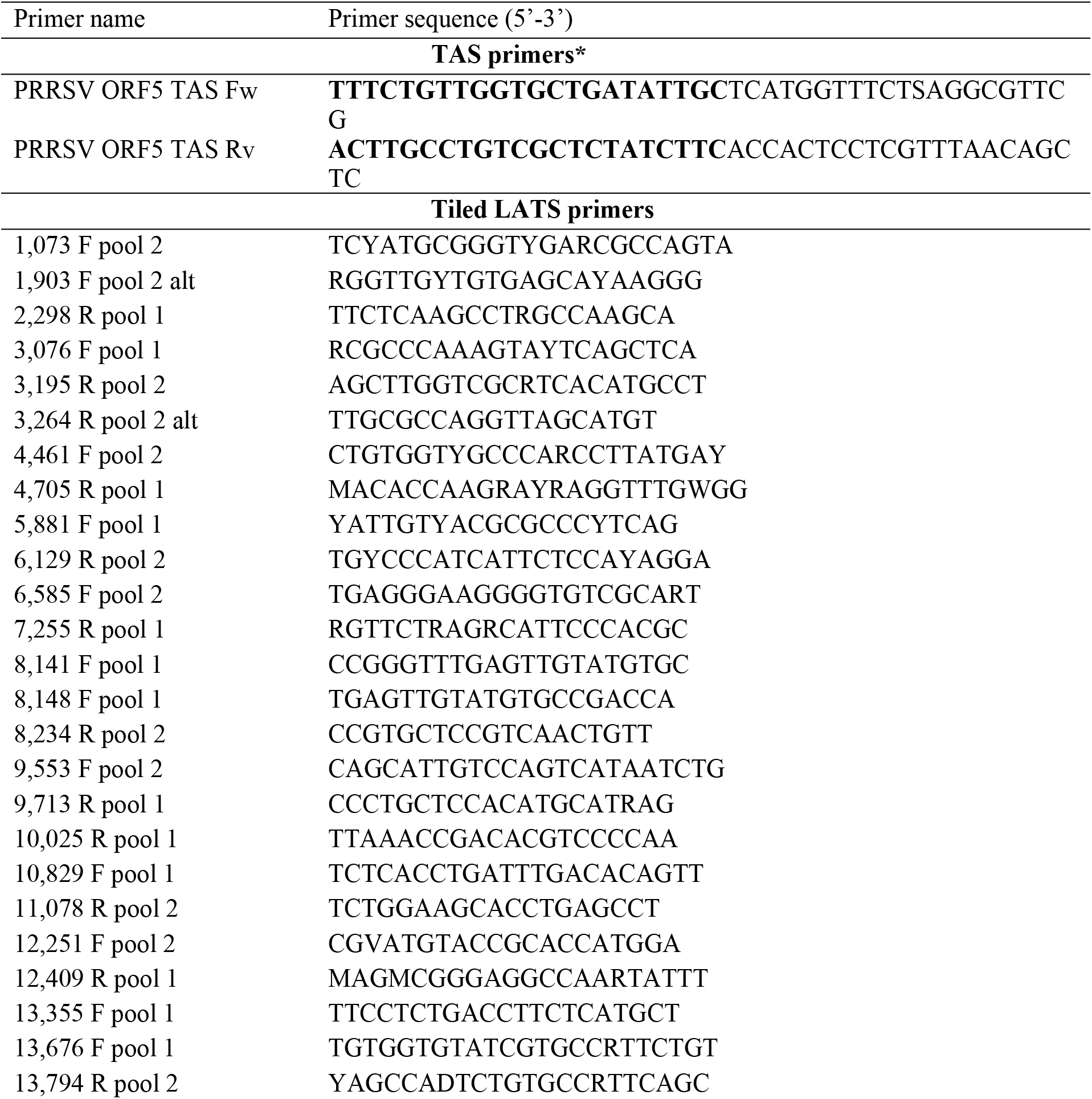
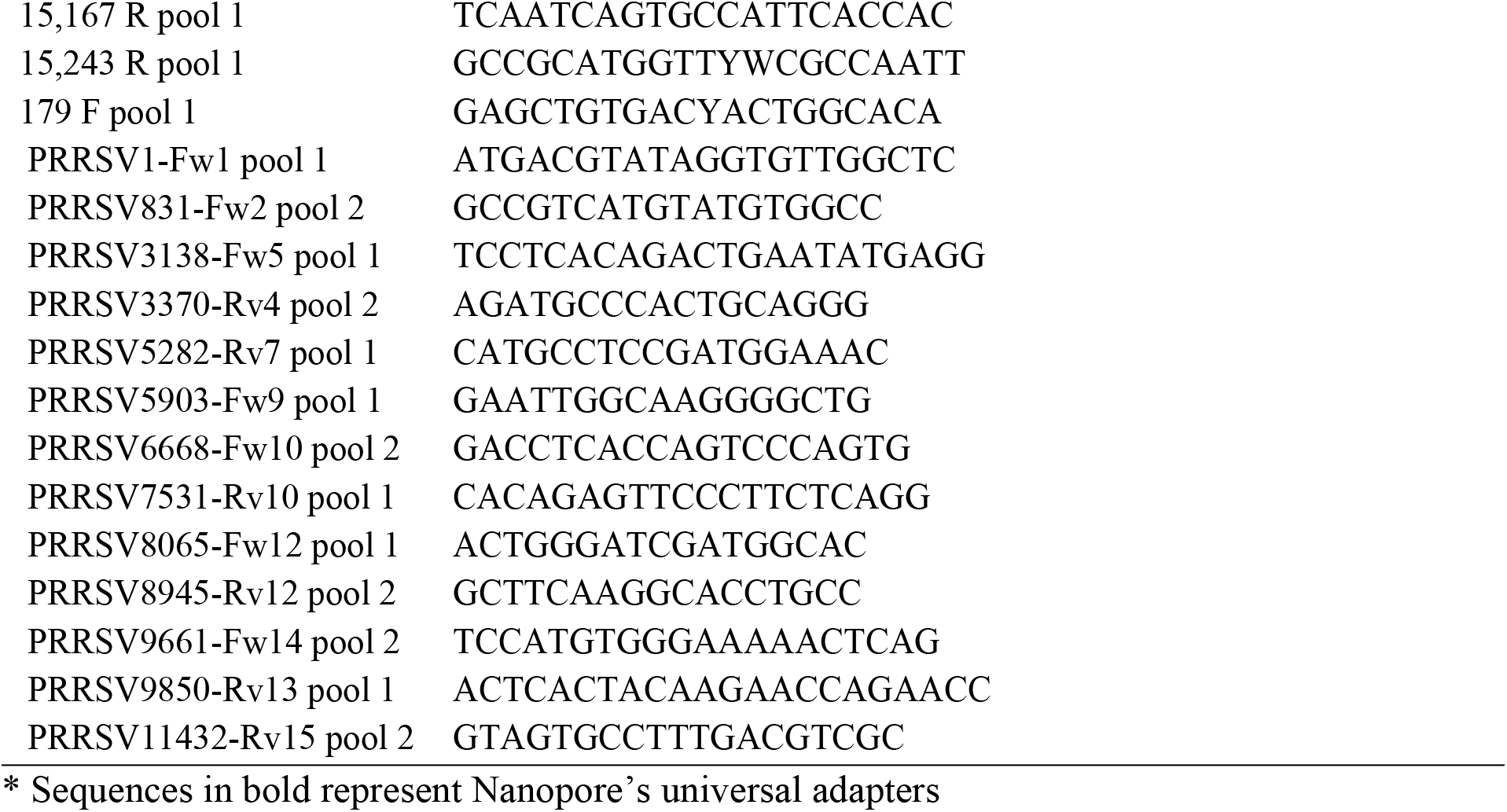
Primers used for TAS and LATS.

### Library preparation for Long Amplicon Tiling Sequencing (LATS)

A multiplex PCR was developed following the ARTIC Network amplicon-based approach for sequencing SARS-CoV-2 (https://artic.network/ncov-2019). Forty-one custom primers were designed manually, targeting approximately 1500bp products with 100bp overlap between different amplicons, based on an alignment of 87 complete PRRSV-2 genomes obtained from GenBank (Figure 2). Primers are provided in Table 1. Libraries were generated using the Native Barcode Kit, EXP-NBD196 and Ligation Sequencing Kit, SQK-SQK109 (Oxford Nanopore Technologies) multiplexing up to 24 samples per sequencing run. Libraries were run on R9.4 flow cells for 6 hours.

**Figure 2.**
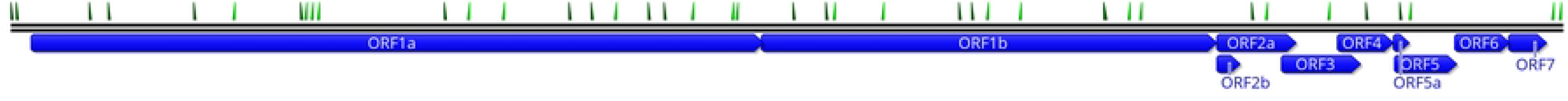
Schematic view of PRRSV genome showing the locations of LATS primer-binding regions.

### TAS analytical sensitivity

The operating range of MinION TAS was determined by sequencing six 10-fold serial dilutions (10^-1^ to 10^-6^) of two PRRSV-1 (strains 03-15 and SD14003) and two PRRSV-2 (strains 1-7-4 and NADC20) cell culture isolates. The serial dilution was initially tested by real-time PCR (RT-qPCR) using EZ-PRRSV MPX 4.0 Master Mix and Enzyme (Tetracore Inc.) and the sensitivity of TAS was compared to this commercial PRRSV RT-qPCR kit. The processes of serial dilution, extraction, library preparation and sequencing were repeated two times independently (experiment 1 and experiment 2).

### MiSeq sequencing comparison

Complete genome sequences of two PRRSV-1 (strains 03-15 and SD14003) and two PRRSV-2 (strains 1-7-4 and NADC20) were obtained using the Illumina MiSeq platform. Briefly, viral RNA was extracted, and cDNA was prepared using SuperScript IV Reverse Transcriptase (Thermo Fisher) followed by second strand synthesis using NEBNext Ultra II Non-Directional RNA Second Strand Synthesis Module (NEB). Libraries were prepared according to the Nextera XT (Illumina) library preparation protocol and then sequenced using the Illumina MiSeq platform.

### Bioinformatic analysis

Raw reads were basecalled and demultiplexed automatically in the MinIT device (ONT). Reads were filtered by quality and minimum size (-q 7 and −l 200) using Nanofilt (27) and then aligned to a PRRSV-2 reference genome, MN073092.1, using the mini_align utility of Pomoxis (https://github.com/nanoporetech/pomoxis). If complete coverage was not obtained, the draft genome was submitted to BLAST and the best hit was used as reference for a new alignment. For the LATS, viral genome was considered for whole genome analysis if a minimum depth of coverage of 20X was obtained for at least 14kb genome length. Consensus sequences were generated using a combination of Medaka (ONT), SAMtools and BCFtools (28). (29) utilities. The commands used can be found in the Supporting information S2. Sequences were aligned using MAFFT (30) and phylogenetic trees were constructed using FastTree (31) with GTR model. For the LATS, same bioinformatic procedures were performed

## Results

### TAS analytical sensitivity

Six 10-fold serial dilutions (10^-1^ to 10^-6^) were used to compare the sensitivity of TAS against RT-qPCR in each of the six dilutions (Table 2). A sequence identity of 100% was obtained within the first 30 minutes of sequencing for the most sample dilutions for PRRSV-1 isolates in both experiments 1 and 2 (Table 2). For PRRSV-2 isolates, a sequence identity of 99.87%-99.93% was obtained for the 10^-6^ dilution in both experiments within the first 30 minutes of sequencing. Two different PRRSV strains of both genotypes were successfully amplified, confirming the specificity and sensitivity of the tailed primers. Noteworthy, although dilutions 10^-6^ resulted in RT-qPCR Cts as high as 32.76-37.36, sequence identities higher than 99.8% were obtained from this dilution across all four isolates in 5 minutes. These results indicate high accuracy, repeatability and broad specificity of the TAS assay encompassing both PRRSV-1 and PRRSV-2 species.

**Table 2.**
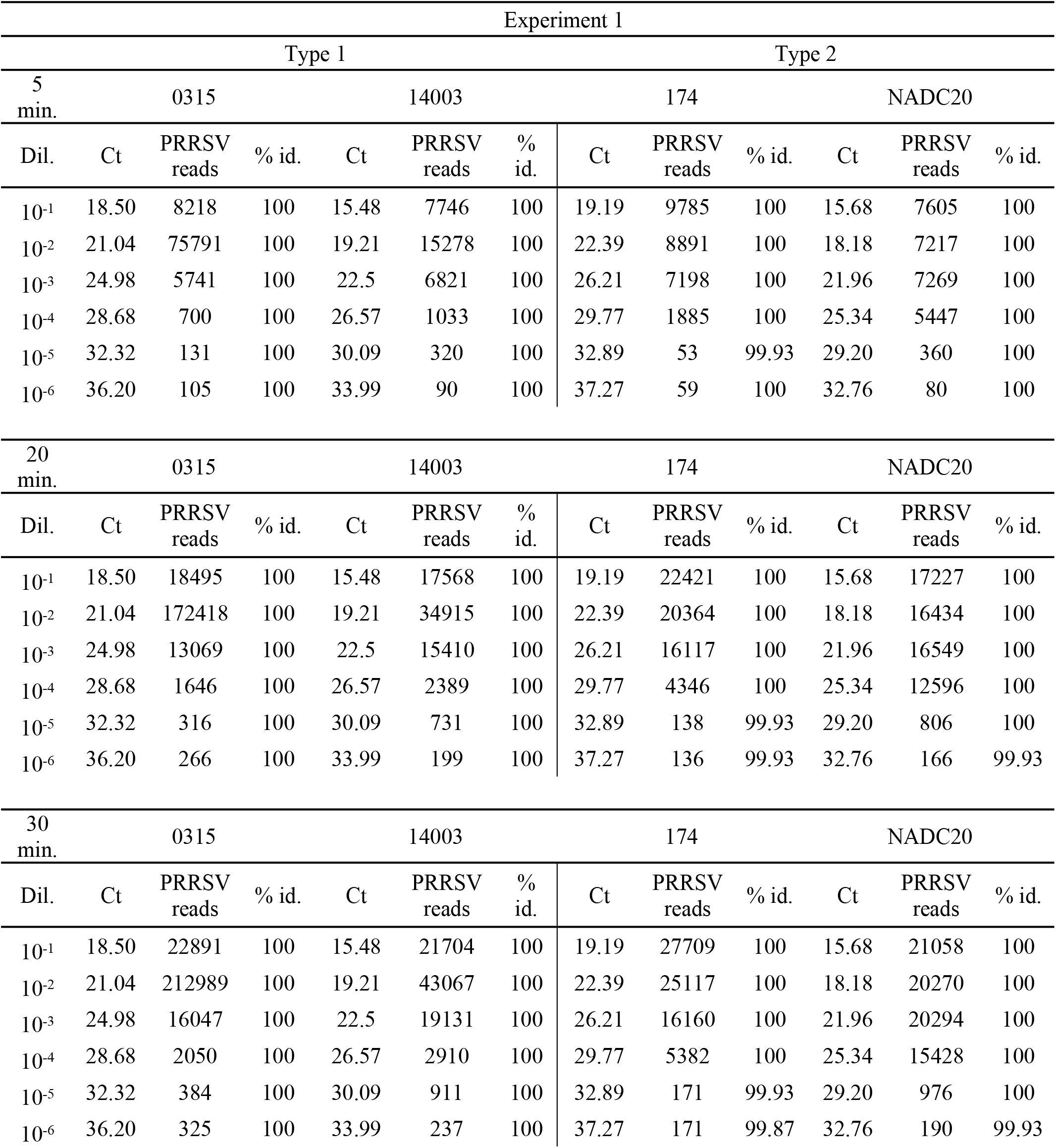
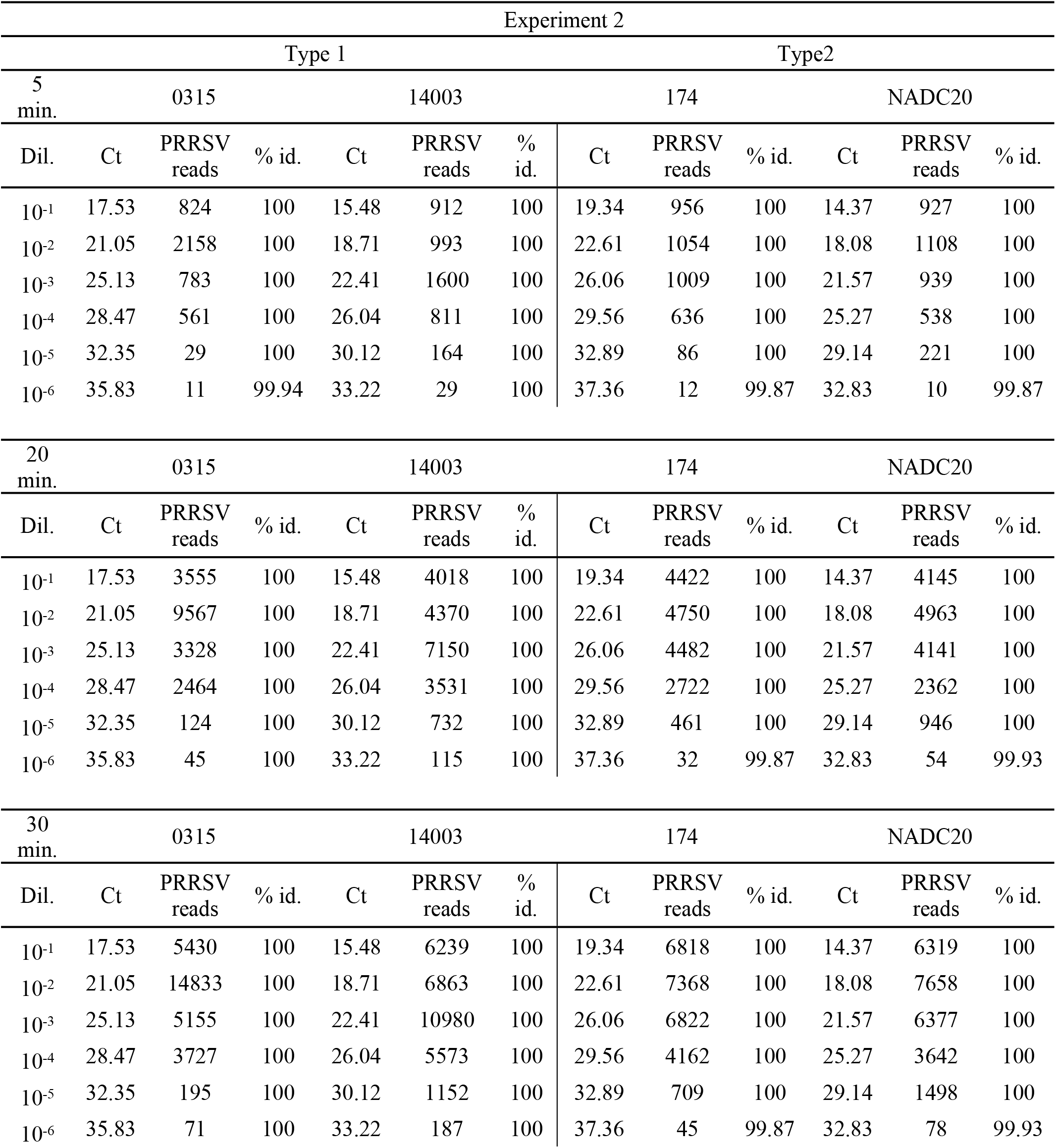
Analytical sensitivity of TAS. Accuracy of consensus sequence from serial dilutions of PRRSV-1 and PRRSV-2 isolates. Two independent experiments were performed on different replicates of the same isolates. A RT-PCR of each dilution was performed to compare Ct values to the number of reads and consensus accuracy.

### TAS on clinical samples

MinION libraries were generated directly from 134 clinical samples (60 sera, 35 oral fluids and 39 processing fluid samples) using the TAS assay and consensus sequences of the full-length amplicon were obtained from all samples. After 1 hour of sequencing, TAS enabled the classification of viruses into lineages 1, 5 and 8 (Figure 3). Full-length amplicon sequences with an average sequencing depth of 421 reads per nucleotide position were obtained from all samples tested.

**Figure 3.**
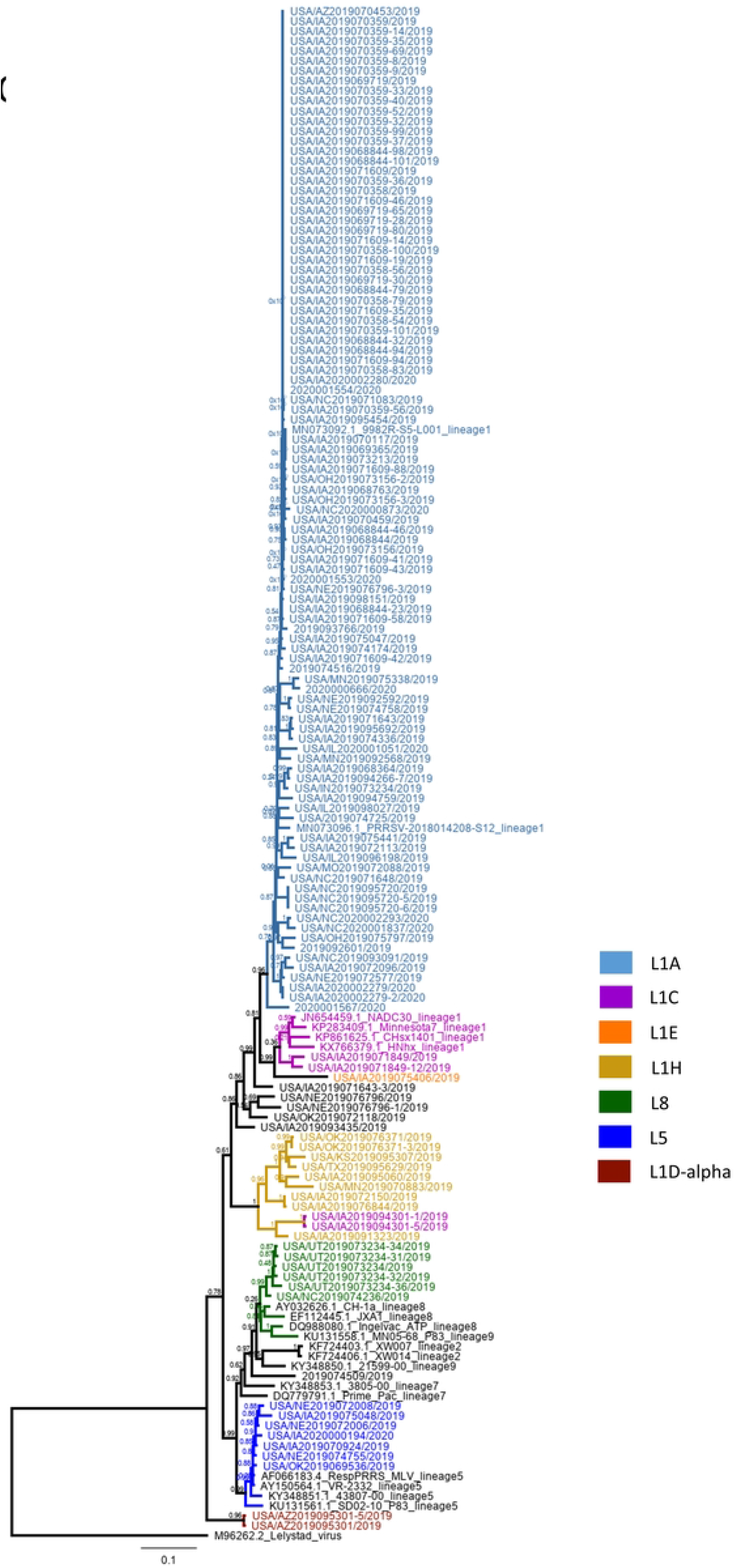
Phylogenetic tree based on partial sequences, spanning ORF4 to ORF6.

### LATS on clinical samples

The utility of the long amplicon tiling sequencing (LATS) on samples from PRRSV outbreaks was assessed by generating libraries directly from lung, serum, oral fluid and processing fluid samples, from a total of 154 field samples. Ninety-two whole genome sequences were obtained (59.7% of clinical samples) and phylogenetic analysis was performed, enabling the classification of these sequences predominantly into lineage 1 and its sublineages, with one sample being classified into lineage 5 (Figure 4). Overall, genome coverage was variable for samples with Ct values higher than 27 (Figures 5 and 6). At least 80% genome coverage was obtained at >20X depth per nucleotide position for 22 out of 35 oral fluid samples (62.8%), 26 out of 39 processing fluid (66.6%), 50 out of 60 sera (83.3%) and eighteen out of 20 lungs (90%). Only 6 serum samples had less than 60% of the genome covered at 20X depth (Figure 5). Most of the samples with RT-PCR Ct values below 25 resulted in nearly complete genome coverage from all specimens tested (serum, processing fluid, oral fluid and lung) (Figure 6). Previously described deletion patterns in NSP2 were observed. Seventy-three whole genome sequences obtained in this study exhibited a 100-aa deletion in relation to VR-2332 reference strain (characteristic of NADC34-like strains), while strain USA/IA2019072096/2019 exhibited a “111+1+19” deletion pattern (characteristic of NADC30-like strains). Sequences USA/NC2019093091/2019 and USA/NC2019095720/2019 contained a 13-aa deletion besides the 100-aa deletion.

**Figure 4.**
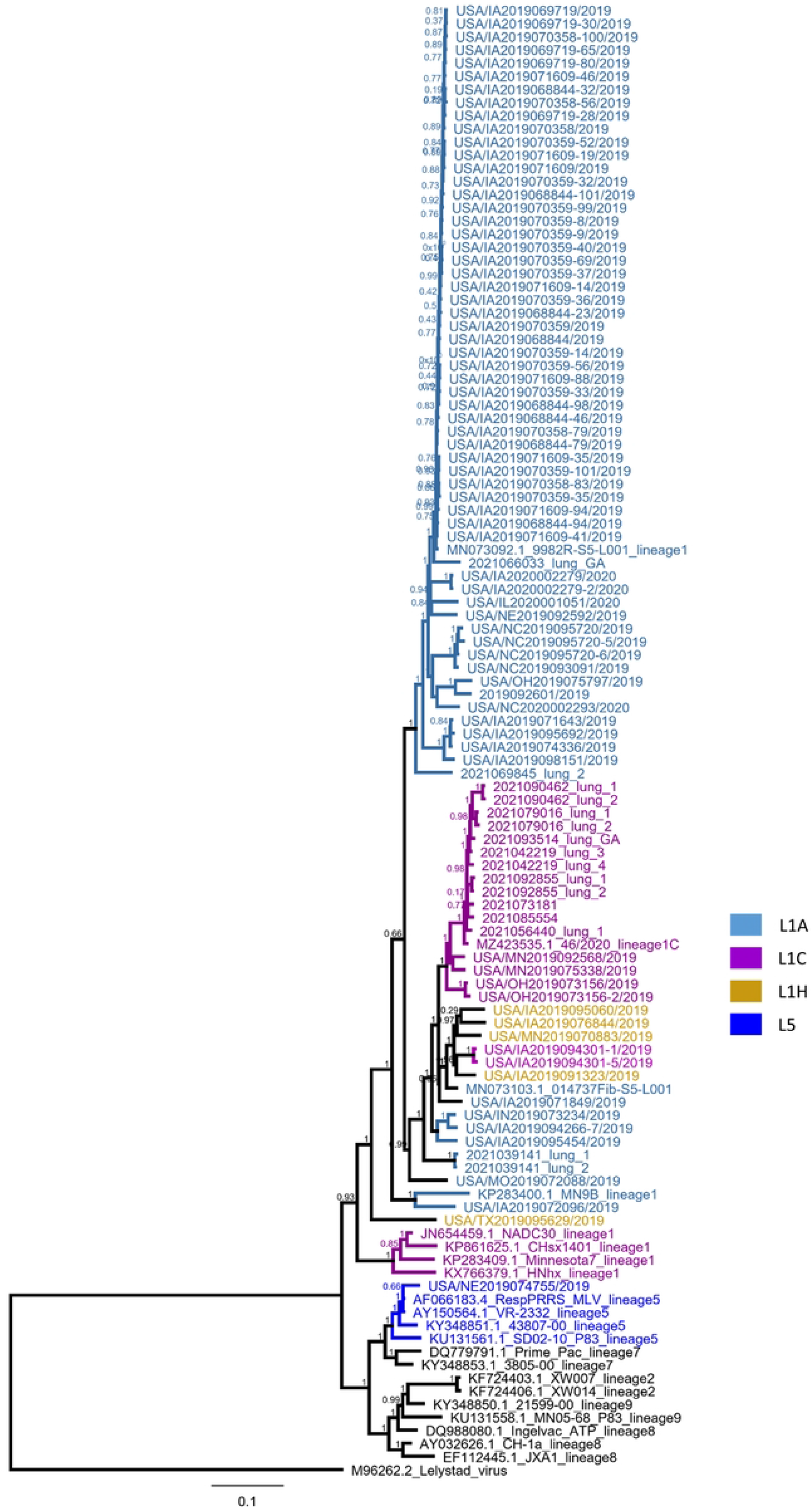
Phylogenetic tree based on complete genome sequences. The type 1 Lelystad virus strain was used as an outgroup.

**Figure 5.**
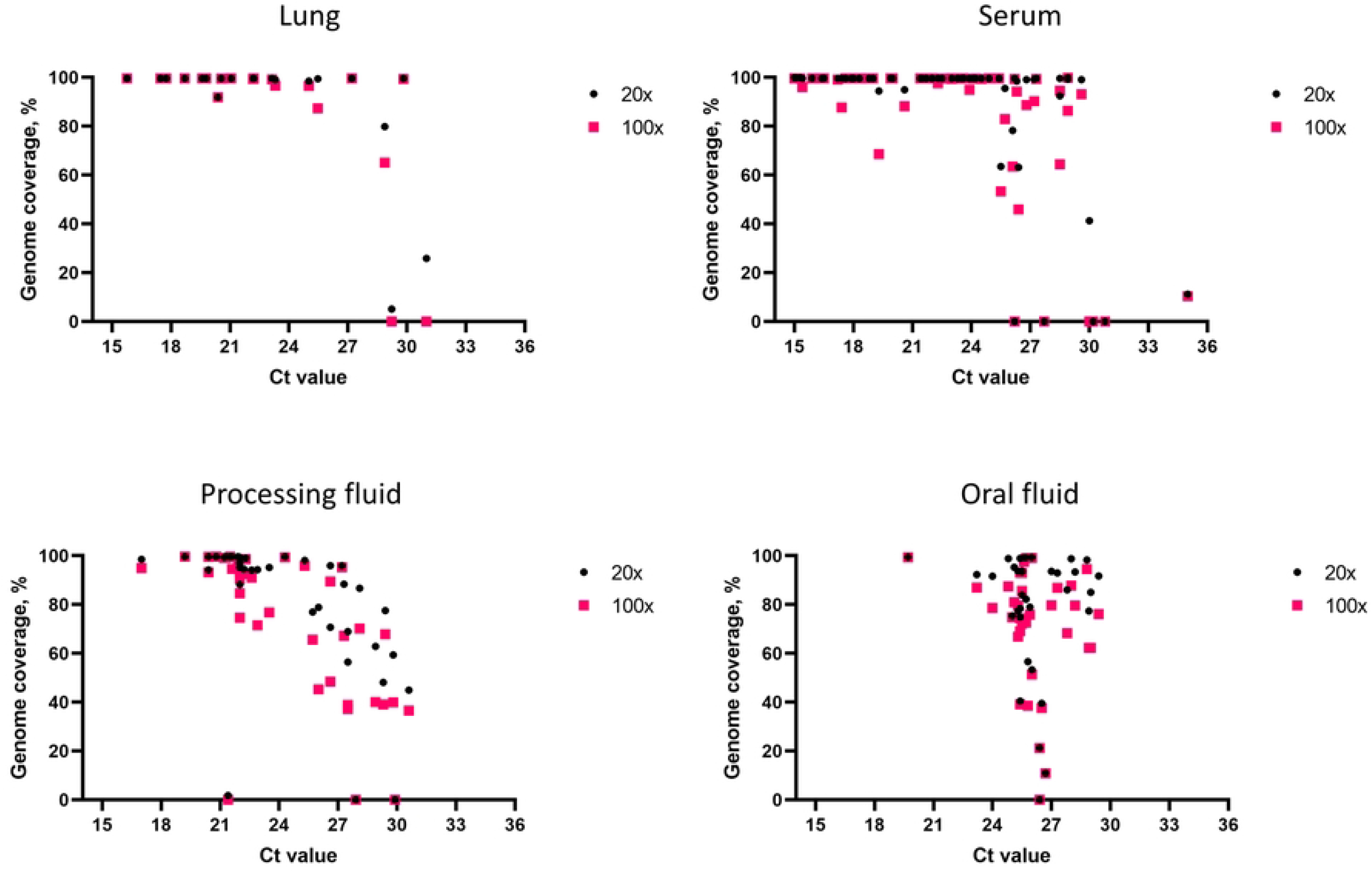
Whole genome coverage at >20X and >100X sequencing depth. Percentage of PRRSV genomes covered by the LATS protocol for lung (n=20), serum (n=60), processing fluid (n=39) and oral fluid (n=35) clinical samples.

**Figure 6.**
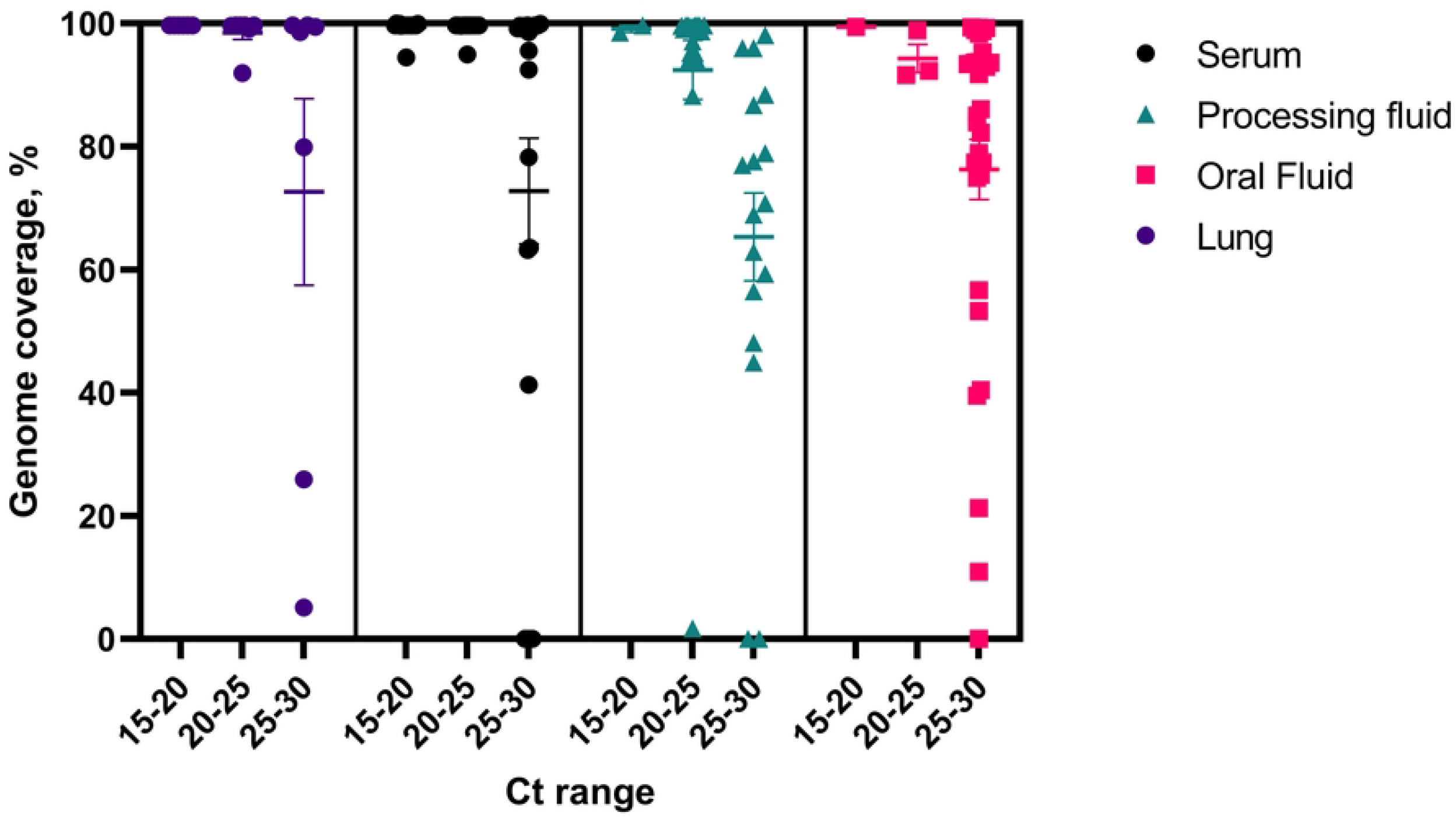
Percent genome coverage per range of Ct value.

### Coverage of ORFs

We also analyzed the coverage for each ORF (ORF1a, ORF1b, ORF2a, ORF3, ORF4, ORF5, ORF6 and ORF7) within the PRRSV genome and compared the depth coverage obtained per nucleotide position across the four samples types included in our study (lung, serum, processing fluid and oral fluid). The percent coverage of each coding region of PRRSV genome was plotted (Figure 7). The coverage of ORF1a, ORF1b and ORF2a presented a similar pattern to the coverage obtained for the whole genome, with good genome coverage in samples with Ct values below 25 and a notable drop in coverage for samples with Ct >27. Lung and serum samples presented the highest overall coverage of ORF1a, ORF1b and ORF2a, whereas a marked drop in coverage in these regions was observed in processing- and oral fluid samples (Figure 7). Notably, a significant increase in sequence coverage was observed in ORF3 to ORF7 which represent ORFs that are transcribed as sub-genomic RNAs from the PRRSV genome.

**Figure 7.**
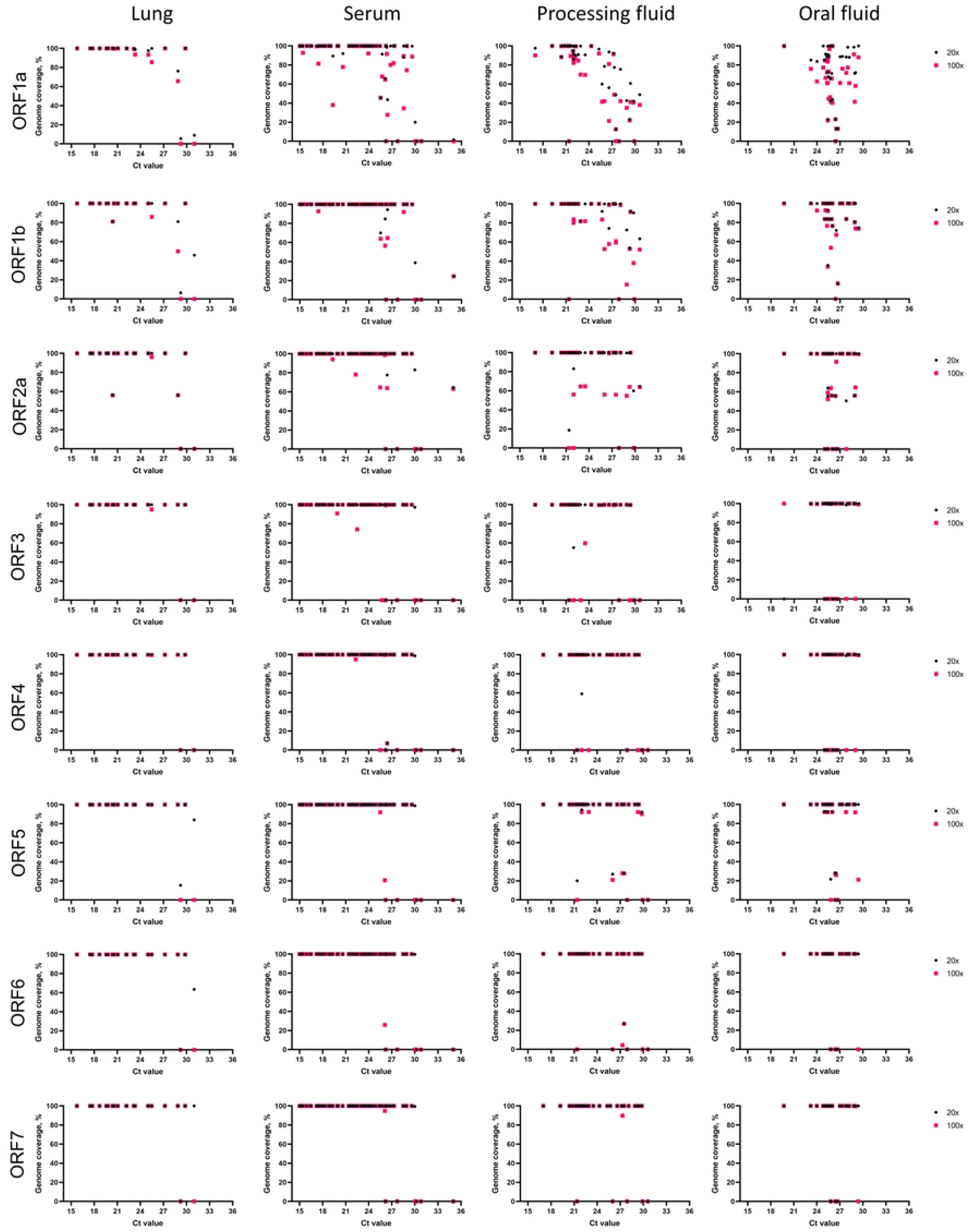
Percent coverage of PRRSV ORFs at >20X and >100X depth by the LATS protocol for lung, serum, processing fluid and oral fluid clinical samples.

### Coverage per time of sequencing

To determine the time points when full genome coverage was obtained after starting the sequencing run, raw reads from serum samples were extracted after 0.5h, 1h, 1.5h, 3h, 4.5h, 6h and were processed as described before. Full genomes with at least 20X sequencing depth were obtained within the first hour of sequencing for samples with Ct values ranging from 15 to 24.9 and >90% coverage was achieved within 4.5h for samples with Ct values of 26.3 and 28.5 (Figure 8). One sample differs from the others, achieving full coverage in 30 minutes despite the Ct of 28.9. At 100X depth, results were more variable and full coverage was obtained within 4.5h for Ct <26.3 and one sample with Ct of 28.9.

**Figure 8.**
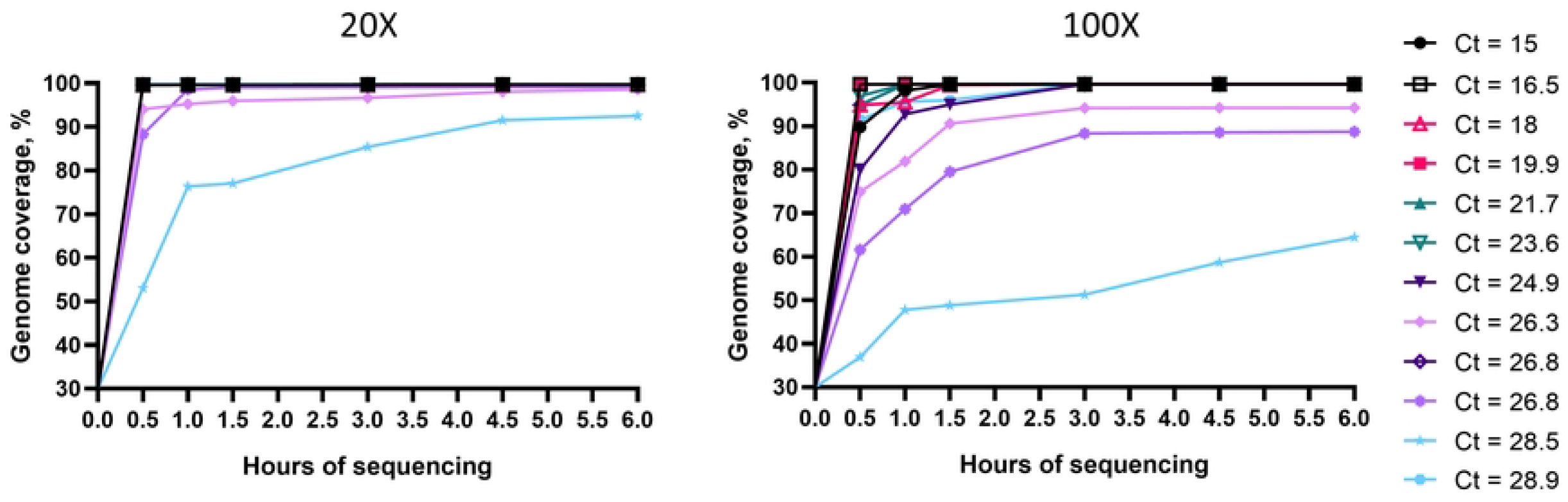
Percent genome coverage per time of sequencing

The average hands-on time of LATS protocol for processing 24 samples from first-strand synthesis to load the library on the flow cell was 7 hours, with a total of 12 hours including the PCR cycling time.

## Discussion

In this study we describe the successful application of two sequencing approaches using MinION platform for accurate PRRSV genotyping directly from clinical samples with a wide range of viral load. These protocols could be useful in all swine-producing regions of the world, especially in developing countries due to the portability, low initial cost and possibility of analysis in real time. The LATS protocol in special, may also be useful for research and surveillance purposes, tracking rapidly-evolving lineages without the need of virus isolation.

The polycystronic nature of the PRRSV genome, which involved ORF2-ORF7, may facilitate sequencing as these ORFs are transcribed as sub-genomic RNAs and are produced in higher abundance when compared to ORF1a and ORF1b during PRRSV infection. The region encompassing these ORFs in the 3’-end of the genome are important for classification of strains and for analyzing interlineage recombination events. ORF7 is also used for identification of strains (32–34) and is fully sequenced by our LATS approach in the vast majority of samples. Thus the LATS developed here could be useful even for the classification of high Ct samples in the impossibility of obtaining whole genome coverage. Use of the LATS procedures developed here also allowed the detection of different deletion patterns in NSP2 region of ORF1a that were previously reported (17,35,36). These results indicate that our primers efficiently amplified this highly variable region. A deeper understanding of NSP2 polymorphism patterns is important for an enhanced classification of lineages, complementing the ORF5-based classification.

The selection of samples originating from different states of U.S. provides more confidence to our phylogenetic classification, which is in accordance with previous studies (35,36). The classification results obtained here show a larger prevalence of lineage 1 and its sublineages, followed by lineage 5 and 8, in accordance with previous reports showing lineages 1, 5 and 8 as the most prevalent lineages inU.S.and China (ref). The viability of this set of primers for sequencing strains of these three lineages suggests that the LATS protocol is suitable for use in responses to PRRSV outbreaks in US, China and other countries where these lineages are present. Lineage classification is based mostly in ORF5 sequences, thus, some whole genome sequences belonging to the same sublineage do not cluster together in the same node in the phylogenetic tree (Figure 4). This could be due to genome recombination. Although PRRSV-2 is the most prevalent in the U.S and China, PRRSV-1 also occurs in these countries and the same LATS approach using a set of primers designed for PRRSV-1 would be useful in outbreaks involving this genotype.

Whole-genome sequencing is highly dependent on sample quality, which could be negatively affected by the action of nucleases, present in the sample, which may vary with the sample type. This is a possible explanation for the variability found in the whole genome coverage analysis (Figures 4 and 5), where it is possible to note that some samples with lower RT-PCR Ct values did not yield full genome coverage when compared to samples with higher Ct values. Differences in sample quality and in the amount and activity of nucleases could have contributed to potential degradation of viral RNA. However, since MinION sequencing allows larger amplicon sizes, a partial genome would provide enough information to enable genotype characterization.

The sensitivity of the TAS was comparable to a commercial RT-qPCR targeting both PRRSV genotypes. However, as RT-qPCR assays only provide positive or negative results, TAS can be more informative since it provides accurate genotyping, revealing mutations at high read depth. TAS primers were designed against the conserved genome regions, which will likely enable long term specific binding to the target regions.

While TAS allows the classification of PRRSV into both genotypes, the assay is more time-consuming than LATS. Taking into consideration the variable nature of PRRSV genome and that PRRSV-2 is the most prevalent and largely spread throughout the U.S, the LATS assay provides more value to the control of this virus in the U.S and other countries where this genotype is highly prevalent. Another advantage of LATS when compared to the TAS procedure is that there is no need for a PCR-barcoding step and PCR clean-up, which decreased the turn-around time. The need for PCR-barcoding in the TAS process also leads to a higher chance of barcode cross-talk, which could lead to wrong barcode assignment in multiplex experiments (37). However, the barcode cross-talk issue can be addressed by further refinements in the bioinformatics pipelines (21). In spite of these drawbacks, the same pair of primers can be used without the adapter tail in a native barcoding library preparation, following very similar downstream steps used in the LATS protocol, thus reducing the turnaround time and chances of contamination. The performance of TAS on high Ct clinical samples depends on sample quality and an efficient first-round PCR. Clinical samples with high Ct values may not have the same sequencing performance observed when sequencing isolates. The same tailed-primers approach was used previously for other viruses such as Newcastle disease virus, reaching an identity of 98.37% compared to the expected consensus within 7 minutes (23) and also for human enterovirus, reaching an average identity >99% when comparing to the expected consensus (21)

The accuracy of MinION sequencing is still affected by its higher read error rate compared to other sequencing technologies, especially in homopolymeric regions (38). Despite this drawback, we obtained overall consensus identities >99% and the quality was sufficient for lineage-level classification of partial and complete genomes. Future development of a robust and user-friendly bioinformatic pipelines for the generation of final data will allow the use of our approaches in a larger scale.

The LATS strategy is effective at generating full genome sequences from samples with Ct values up to 30, with a turnaround time of 11 hours from RNA to final DNA library. This protocol offers a cost-effective and informative system that can be used in research and diagnostic laboratories and in the field, providing invaluable information for veterinarians, and epidemiologists working on the control of PRRSV. As shown by the rapid shift of prevalence from lineage 9 to lineage 1 within a 5-year frame, whole genome sequence classification complementary to ORF5 is essential.

The rapid turnaround time, portability and ease of use offer significant advantages over traditional diagnostic approaches or other sequencing platforms. The protocols presented here may become valuable tools with potential for field applications during PRRSV elimination programs.

## Data Availability Statement

The sequence dataset used here is available in GenBank under the accession numbers MW592708-MW592739, MW810509-MW810586 and OQ361807-OQ361824. Raw reads are available in the NCBI Sequence Read Archive (SRA) under BioProject number PRJNA934299

## Funding

The work was funded by National Pork Board (project #18-176, DGD). Sequencing infrastructure established at the Cornell Animal Health Diagnostic Center Virology Laboratory was funded in part by the USDA Animal and Plant Health Inspection Service (agreement no. AP20VSD&B000C020, DGD). The funders had no role in study design, data collection and analysis, decision to publish, or preparation of the manuscript.

## Conflict of interest

DGD and LCC have filed a provisional patent application on the PRRSV TAS and LATS assays developed in this study (US serial no. 63/413,744).

## Supporting information

**S1 Table**. **Description of clinical samples used in this study and respective GenBank accession numbers.**

**S2 Table. Bioinformatic commands used for generating consensus sequences.**

